# Effect of the microbiome on pathogen susceptibility across four Drosophilidae species

**DOI:** 10.1101/2025.10.22.683894

**Authors:** Hongbo Sun, Ben Longdon, Ben Raymond

**Affiliations:** Centre for Ecology & Conservation, Faculty of Environment, Science, and Economy, University of Exeter, Penryn Campus, Penryn, United Kingdom

**Keywords:** Microbiome, Drosophila, Bacterial infection, Viral infection, DCV, *Providencia rettgeri*, *Staphylococcus aureus*

## Abstract

The microbiota has been shown to play an important role in host susceptibility to infections in some hosts. However, less is known about whether microbiota-mediated effects are consistent across host species, as our understanding of such interactions may be affected by publication bias. Following on from a large study of 36 species of Drosophilidae challenged with four bacterial pathogens, we identified two candidate host species that might have protective microbiome based on low susceptibility and high abundance of the culturable microbiota; we selected two other host species for comparison. We tested whether germ-reduced flies, and flies with a natural or re-constituted microbiome, varied in their susceptibility to systemic infection with two bacterial pathogens (*Providencia rettgeri* and *Staphylococcus aureus*) and one viral pathogen (Drosophila C Virus). The composition and abundance of the bacterial microbiota varied between species and microbiome treatments. We found an overall interaction between host species and pathogen type, confirming previous work that host species vary in their susceptibility in a pathogen-specific manner. Similarly, we found that microbiome treatments had differing effects on host survival among host species, although effect sizes tended to be small. In *D. putrida* individuals with manipulated microbiomes showed increased susceptibility to all pathogens tested; in other hosts altered susceptibility was pathogen dependent. While there are always challenges to manipulating microbiomes, especially across multiple host species, our results indicate that host microbiota may play limited roles on survival in systemic infection in these four species. This work demonstrates that caution is required when generalising about the potential beneficial impact of microbiomes.

## Introduction

Animals and plants survive and evolve in association with microbes [1]. The microbiome is the collection of microorganisms (bacteria, fungi, and viruses) that inhabit a particular host [2]. Host-microbe interactions span from mutually beneficial to pathogenetic [3, 4]. Microbiome composition varies across host species [5], and can contribute to different aspects of host physiology, such as development, nutrition and immunity [3, 6–8]. Although many microbes play vital roles in host physiology, the microbiome can still have pathological effects [9]. Hosts use various strategies, such as stomach acidity, mucosal barriers, and immune response, to shape and regulate their microbial communities [9–13]. However, excessive immune response can have physiological costs, leading to rise of inflammation and weakened overall host defence [9]. Establishing tightly regulated symbiotic relationships typically requires extensive anatomical specialisation and millions of years of coevolution [14]. These conditions are not met for the majority of microbiome members. However, microorganisms can have commensal relationships with hosts without having undergone natural selection to benefit hosts [15]. Thus, the observation of phylosymbiosis, where ecological relatedness of host-associated microbial communities parallels the phylogeny of related host species [5, 16–20], may not necessarily imply that the microbiome has coevolved with hosts [15]. In addition, we are often too ready to infer microbiome functions based solely on sequencing data and associations with host traits, without considering the underlying ecological complexity [21].

For example, the microbiome can have positive, negative or neutral effects on a host’s susceptibility to infection, and such effects can be context-dependent [22]. There are a number of examples where the microbiome can provide protection against infection in particular host species [23–25]. The human gut microbiota protects against colonisation by disease-causing bacterial pathogens [24]. Such colonisation resistance can be mediated by “nutrient-blocking”, driven by either high microbiome diversity and the presence of certain key microbiome taxa [26]. In woody plants, the rhizosphere and endophytic microbiome are linked with certain pathogen defences [10, 25]. This involves the selection and recruitment of beneficial microbes in the rhizosphere [27], and secretion of defensive enzymes [28], phytohormones [29], and secondary metabolites [29, 30] from the endosphere microbiome.

However, in some cases, the microbiome can facilitate infection [31, 32]. In some cases, normally harmless commensals may become pathogenic in immunocompromised hosts [33]. Prolonged microbiome dysbiosis can lead to the accumulation of pathobionts, the loss of both key commensals and the loss of the taxa diversity, which comes along with the dysregulated immune defences [34]. Microbes that have pathogenic potential within certain contexts are termed “pathobionts” [31, 32, 35, 36]. In addition, not all microbial interactions alter infection susceptibility; some microbes coexist without impacting pathogen resistance [37], while others exert effects that are highly strain-dependent [38].

Invertebrates represent more than 90% of all living animal species [39]. The microbiomes of simple invertebrates such as nematodes, bees, and flies display both shared and species-specific characteristics [14]. Compared to vertebrates, whose microbiomes are typically dominated by Bacteroidetes and Firmicutes, invertebrate microbiomes tend to harbour fewer microbial species and are mainly dominated by Proteobacteria and Firmicutes [14]. Microbial composition can differ across species: wax moth larvae, *Galleria mellonella* are primarily colonised by *Enterococcus* which can stimulate host immunity to increase protection from Gram-positive pathogens [40]. In nematodes, wild *C. elegans* collected from their natural habitats exhibited extensive microbial diversity. Over 250 bacterial genera were identified in rotting apples, which were mainly dominated by Enterobacteriaceae and acid-producing bacteria [41], a composition similar to *D. melanogaster* [42, 43]. Both host-specific factors, such as immune factors (e.g., antimicrobial peptides, PGRPs, DUOX), gut structure, nutrition; and environmental factors, such as habitat, diet, and microbial exposure, shape invertebrate microbiota [44–47]. These microbial communities can also modulate host defence in a context-dependent manner [48–52]. For instance, in the mosquito *Aedes aegypti,* the microbiome suppresses viral replication and transmission by manipulating reproduction or direct pathogen-blocking [53]. While in honey bees, the gut microbiome aids in clearing *S. marcescens* infections through both antimicrobial peptide mediated pathogen clearance and colonization resistance [54].

*Drosophila melanogaster* has been well-developed as a model for microbiome research, with the gut structure similar to that of vertebrates and a relatively simple and easy-to-manipulate microbiome [42, 55]. Data on wild-caught flies have shown that 85% of the natural bacterial microbiome is composed of only three bacterial families, including Enterobacteriaceae, Acetobacteraceae and Lactobacillales, despite the phylogenetic, ecological, and geographical diversity of the hosts [56]. Laboratory-reared flies showed that the intestinal bacterial microbiome represents only a small fraction of the environmental microbial communities, suggesting that Drosophila’s low-diversity communities might be linked to the host filtering effects [56, 57], although diet can contribute to bacterial microbiome composition [56, 58]. However, there is still variation between host species in the microbiome composition at lower taxonomic levels [56, 59].

Many studies have revealed the importance of the microbiome of *D. melanogaster* on host fitness, including its role during infection [6, 48, 56, 57, 60–67]. In *D. melanogaster*, *Lactiplantibacillus plantarum* and *Acetobacter tropicalis* enhance host survival and decrease pathogen loads during oral infections of *P. entomophila* [48]. These protective effects occur within the gut lumen, where *L. plantarum* acidifies the environment to suppress harmful pathogens, while *A. tropicalis* counters potential adverse effects by neutralizing acid, thereby maintaining gut homeostasis [48]. Microbiota-mediated resilience mechanisms are also necessary in stabilising intestinal homeostasis during oral infection [68]. In addition to influencing pathogen colonisation, the microbiota modulate host immunity through mechanisms such as immune priming, where prior exposure to commensals or sublethal infections can enhance the host’s ability to mount a faster or stronger immune response upon subsequent pathogen exposure [69].

Comparative genomic studies have revealed the different evolutionary patterns for immune-related genes among Drosophilidae species [70]. For example, Toll and IMD pathways are known for their role in defence against bacterial pathogens [71]. However, genes involved in these pathways varied significantly across species [70]. Genes encoding effector proteins (for example, AMPs) and recognition proteins are much more likely to vary in copy number across species than genes encoding signalling proteins [70]. Microbiome-mediated immune priming is a process involving an increase in the density of circulating haemocytes and AMPs [72]. In this case, we cannot assume immune priming is common or even generally adaptive across Drosophilidae, let alone all insects. Here, we aimed to explore if intact microbiomes consistently confer protection across multiple species within this family, and how these microbiome-host interactions vary across different pathogens.

Following a study of 36 species of Drosophilidae challenged with four bacterial pathogens, we identified two host species (*Drosophila putrida* and *Scaptodrosophila pattersoni*) that we hypothesized had a protective microbiome based on their low susceptibility in systemic infection and high abundance of the culturable microbiome [73]. We selected two other species as comparators (*Drosophila melanogaster*, *Zaprionous davidi*) which are phylogenetically distantly-related. We examined the susceptibility of these species with either intact or manipulated microbiomes. To manipulate microbiomes, we used antibiotics to produce a “germ-reduced” treatment. We then reconstituted fly microbiomes with either an *Acetobacter* isolate, which is one of the dominant bacterial genus in Drosophilidae microbiomes; or a *Providencia,* which may act to prime the host against related pathogenic bacteria [74]. For each of these four treatments, and for each of the four host species, we infected flies independently with three pathogens: a Gram-negative bacteria, *Providencia rettgeri* [75]; a Gram-positive bacteria, *Staphylococcus aureus* [76, 77]; and an RNA virus, Drosophila C virus (DCV) [76, 78, 79].

Our study tested whether the presence, disruption, or specific constituents of the host microbiota affect host susceptibility from systemic infection. We found that intact microbiomes could either increase or decrease risk of pathogen infection relative to germ-reduced microbiomes and this effect varied with host species and pathogen. Microbiome treatments had a strong effect on overall survival in *D. putrida*, but only weakly affected survival in a few pathogen-species combinations in the other fly species. These effects were linked to overall disruption of the microbiome rather than variation in the abundance and composition of particular bacterial taxa.

## Methods

### Approach

Here we examined the role of bacterial microbiome in determining susceptibility to pathogen infection across four host species (*D. melanogaster*, *D. putrida*, *S. pattersoni*, and Z*. davidi*). We compared survival of these flies across four treatments: i) conventionally reared flies with intact microbiota, ii) ’germ-reduced’ flies that had been treated with antibiotics to reduce the microbiota; or germ-reduced flies reassociated with iii) *Acetobacter* or iv) *Providencia* bacteria. The *Acetobacter* and *Providencia* were isolated from *D. putrida*, which based on previous findings [73] we hypothesised could be providing protection. We elected to use microbes from a single fly species (Table 1) to reconstitute microbiomes, rather than the specific taxa from each host, in order to avoid additional microbe-by-host-by-pathogen three-way interactions in our experimental design. Flies from each species-treatment combination were either inoculated with a control (Ringer solution) or one of three pathogens: *Providencia rettgeri, Staphylococcus aureus*, or Drosophila C virus (DCV). In addition, we collected a small subset of flies from each treatment (prior to infection) for 16S amplicon sequencing and RT-qPCR to assess changes in the bacterial microbiome.

**Table 1:** Bacteria strain, virus isolate, media, OD and source information. . *Providencia. clone JH10* and *Acetobacter. sp* were isolates used for oral infection and reconstitution of fly microbiome and were isolated from *D. putrida*.

| Bacteria and virus strain | Medium | Use | OD<sub>600</sub> in inocula | Source |
| --- | --- | --- | --- | --- |
| *Staphylococcus aureus* PIG1 | LB (Lysogeny Broth) | Pinprick infection | 1.0 | [77] |
| *Providencia rettgeri* Dmel | LB (Lysogeny Broth) | Pinprick infection | 1.0 | [75] |
| *Providencia*. clone JH10 | LB (Lysogeny Broth) | Oral colonisation | 2.0 | this study * |
| *Acetobacter*. sp | MRS | Oral colonisation | 2.0 | this study * |
| DCV | NA | Pinprick infection | NA | [79] |

We treated newly emerged adult flies with an antibiotic cocktail for seven days before rearing them on sterile antibiotic-free media for three days to clear antimicrobials from tissues (Figure 1) to reduce the chance of antibiotic persistence affecting the bacterial pathogen. In prior studies, we compared the microbiome abundance between conventionally reared flies and the germ-reduced flies to confirm that antibiotics can efficiently clear/reduce the culturable host microbiome (Figure S1).

**Figure 1.**
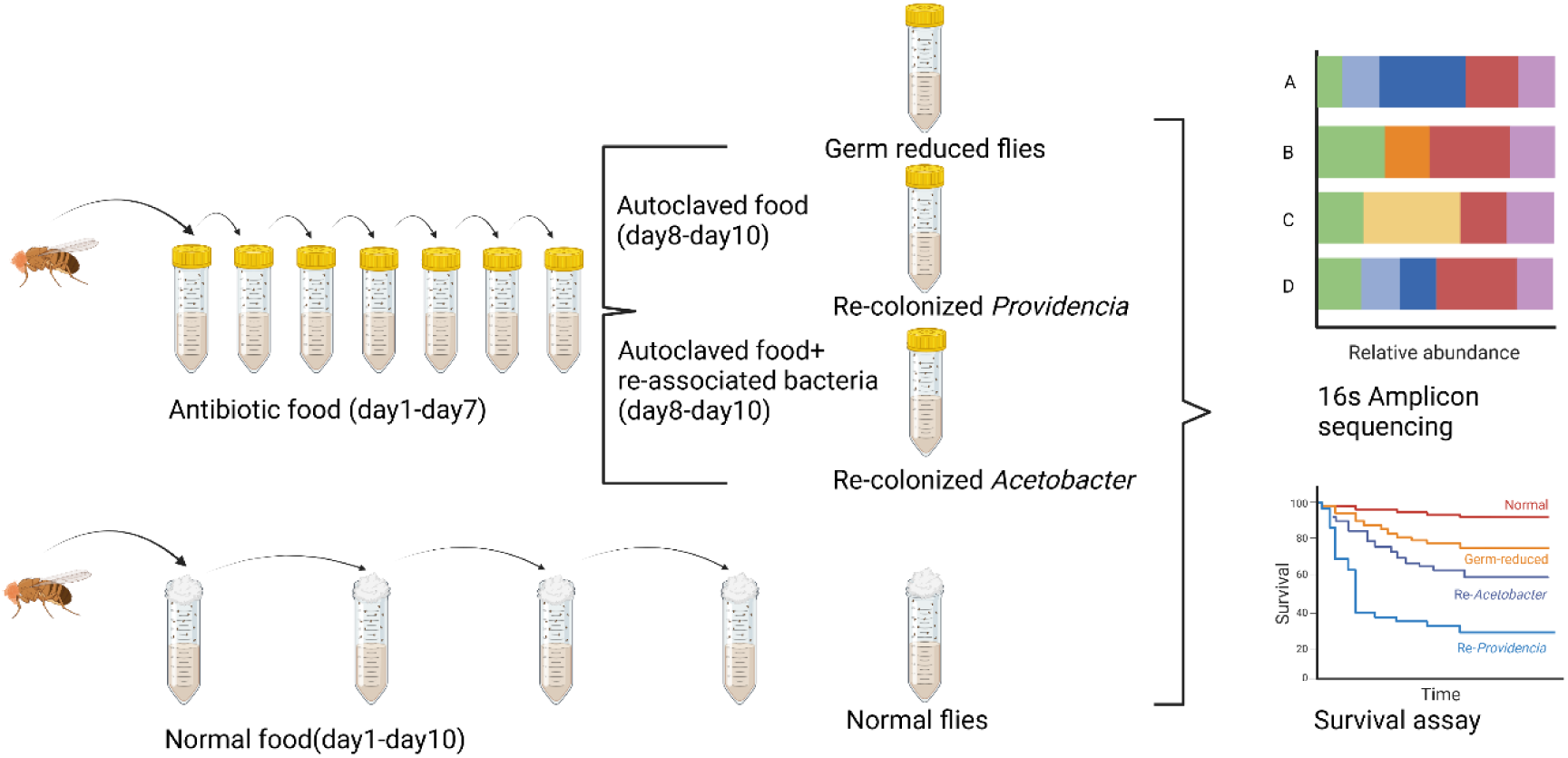
Experimental design for 16s amplicon and survival assay. Normal flies were reared on normal malt food. Germ-reduced flies were reared on autoclaved malt food supplemented with antibiotics. The same *Acetobacter* and *Providencia* isolates were used to reconstitute microbiomes in all fly species. All types of flies were challenged with different pathogens at the same age.

### Pathogen and culture conditions

Bacterial cultures were initiated from frozen glycerol stocks, by plating on agar for 2–3 days. A single colony of bacteria was used to prepare overnight cultures of 5 mL in 30 mL sterile universals (Table 1). Overnight culture time were strictly controlled to no more than 12 hours, with the exception of *Acetobacter. sp* which was cultured no more than 36 hours to standardise physiology. All bacterial cultures were shaken at 180 RPM at 30 ℃. Following overnight growth, stationary phase cultures were diluted with respective culture media (Table 1) to the desired optical density (OD) value before being used in pinprick infections (for pathogen) or oral feeding (to reconstitute the microbiome). DCV isolate used here (DCV-C) was isolated from lab stocks established by wild capture in Charolles, France [79].

### Normal flies, germ-reduced flies and microbe-recolonised flies

All flies were maintained in multi-generation stock bottles (Fisher brand) at 22°C, ∼ 50% relative humidity in a 12-hour light-dark cycle. Each stock bottle contained 50 ml of one of three varieties of *Drosophila* media, which were chosen to optimise rearing conditions (full details in Supplementary Table 1). Male flies were used to avoid any effect of sex or mating status, which has been shown to influence the susceptibility of female flies to various pathogens [80–82]. Newly emerged 0-1day male flies were transferred to vials containing standard malt media, and here after in formal assay, all flies were kept on this media to avoid any effects of food media on the microbiome. Food cooking and dispensing for normal flies was carried out in the normal cooking room, sterile food was cooked and dispensed in a class 2 microbiological safety cabinet.

These flies were then transferred to either fresh malt media (normal flies) every three days for 10 days (age 10–11 days, flip three times before infection), at which point they were inoculated; or fresh autoclaved malt media supplemented with an antibiotic cocktail, where flies were tipped onto fresh antibiotic media every day for seven days. The antibiotic cocktail was comprised of Kanamycin (50 µg /mL), Rifampicin (50 µg /mL), Cefotaxime (10 µg/mL), and Ofloxacin (1µg /mL). After seven-days on antibiotic media, flies were transferred to either sterile malt food (germ-reduced flies) or sterile malt food supplemented with *Providencia JH10* or *Acetobacter. sp* (150 µL cultures standardised to OD_600_ 2.0) for three days (microbiome re-constituted flies), at which they were pricked together with the conventionally reared normal flies (Figure 1).

For oral colonisation, bacterial cultures were inoculated onto the food medium two hours before flies were placed into the respective tubes, to avoid contact between flies and free liquid. 50 mL bioreactor centrifuge tubes (Thermo Scientific, catalogue number: 332260) were used as food containers for both germ-reduced flies and microbiome re-constituted flies. These tubes have a 0.2 μm filter membrane that allows for controlled gas exchange and air circulation without contamination. Standard 50ml Falcon tubes (Thermo Scientific, catalogue number: 339652) plugged with non-absorbent cotton wool were used for normal flies.

### 16S amplicon sequencing

We used the 16S amplicon sequencing method to study the bacterial microbiome structure and relative abundance of taxa within each treatment. We used a 16S rRNA gene metabarcoding approach from extracted RNA (reverse transcribed into cDNA) to select metabolically active bacteria and exclude environmental contaminations [83]. We collected 109 samples (control samples included) covering four types of flies prior to the survival experiments, we removed two samples in our final analysis as the samples appeared to have been swapped by error (7-54-N and 3-54-RP). Each sample was a pool of eight flies. Flies were treated with Trizol reagents and homogenised using a bead beater. Samples were homogenized using the OMNI Bead Ruptor at speed six for 3 × 15-second cycles, with 10- second pauses between each cycle. Direct-zol RNA Miniprep Kits (Zymo Research) were then used to extract total RNA. Nucleic acid-free water blanks were processed with the same nucleic acid extraction kit as a negative control. Bacterial samples in the ZymoBIOMICS Microbial Community Standard kit (Zymo Research) were also processed for RNA extraction as a positive control. Extracted RNA was then normalised to 500ng/µL, and cDNA was synthesised using a GoScript™ Reverse Transcription System (Promega). cDNA products were then quantified on Qubit (ssDNA Assay Kit, Thermo Fisher) followed by normalisation (20 ng/µL). We also used DNA standard samples (ZymoBIOMICS Microbial Community DNA Standard (Zymo Research)) as a template control.

The v3-v4 region of the cDNA 16S rRNA gene was amplified using primer pairs 341F (5’-CCTACGGGNGGCWGCAG-3’) and 785R (5’-GACTACHVGGGTATCTAATCC-3’) [84]. DNA libraries were prepared by PCR cycle of pre-amplification denaturation of 95 °C for 3 mins, 18 cycles of annealing at 95°C for 30 s, 55 °C for 30 s, 72 °C for 30 s, and a final extension at 72 °C for 5 min. Each 25 μL reaction comprised 12.5 μL NEBNext high fidelity PCR master mix (New England Biolabs), 5 μL each 1μM primer, and 2.5 μL template. PCR products were then purified through Agencourt SPRIbeads (Beckman Coulter), which were able to purify 16S amplicon away from free primers and primer dimer species. Illumina sequencing adapters (barcodes) were then ligated using a 10-cycle PCR of the same PCR cycling conditions as described above, with slightly different 25 μL PCR mix (12.5 μL NEBNext high fidelity PCR master mix, 5 μL IDT indexing primer, 7.5 μL template).

The sequencing run was performed on an Illumina MiSeq PE300 system at the University of Exeter sequencing centre. Raw Illumina data were processed using bcl-convert (v4.0.3) Illumina [85] to demultiplex into samples and generate fastq files. Reads were trimmed using Cutadapt Martin [86] (version 4.0) to remove sequencing adapters and low-quality (<Q11) bases from the 3’ end, reads shorter than 150bp were discarded. 16S reads were de-phased using custom scripts. During library preparation, short, variable-length, non-biological sequences were added to the start of the read. This increases sequence diversity and improves sequencing quality. The PCR primers were removed using Cutadapt Martin [86] (version 4.0), so all reads not having the correct primer sequence were discarded.

The resulting sequencing reads were processed using the DADA2 [87] package (version 1.30.0) within R version 4.3.1 (R core team, 2023) in R studio (Posit team, 2024). The forward reads were trimmed to 270 bp, and reverse reads were trimmed to 240 bp using the “filterAndTrim” function based on the sequencing quality. Reads were then dereplicated by the “derepFastq” function, and then applied by the core sequence variant inference algorithm “dada” and merged by the “mergePairs” function. Taxonomy were assigned by the “assignTaxonomy” function based on a DADA2-formatted 16S reference database (Silva version 138.2) [88].

The alpha diversity differences between treatments were tested using a linear model. Beta diversity was assessed through unconstrained principal coordinate analysis (pcoa), based on the bray-curtis dissimilarity matrix at the ASV (amplicon sequence variant) level. The pcoa was performed using the “ordinate” function within the phyloseq package in R [89]. We calculated relative read abundance, which is the proportion of every read belonging to each ASV using a customised R script based on the package dplyr and ggplot2 [90, 91].

### Measuring bacterial load

We measured colony-forming units (CFUs) of bacterial microbiota for individual male flies for normal and germ-reduced flies on both LB agar and MRS agar plates. After one-week antibiotic food treatments, we selected three time points (day 1, day 4, and day 7) in the following week to measure. Anaesthetised flies were transferred to 1.2 mL homogenisation tubes, each containing a 4 mm steel ball and 200 µL of 70% ethanol. The flies were rinsed with 70% ethanol briefly before being homogenised in 200 µL of sterile saline solution. Homogenisation was performed using a Tissue Lyser II set to 30 Hz for three minutes. The homogenates were serially diluted from 10^-1^ to 10^-5^ and plated on LB agar or MRS agar in two technical replicates. The plates were then incubated overnight at 30°C. Bacterial loads were measured as colony-forming units (CFUs) per fly, following the formula:

CFUs per fly = (Colony counts × Dilution factor × Homogenate volume) / Plating volume.

Bacterial relative 16S gene expression for each type of flies were quantified by RT-qPCR method. RT-qPCR was performed for both bacterial *16S rRNA* gene and the housekeeping gene *RPL32* using the Sensifast Lo-Rox SYBR kit (Bioline) on an Applied Biosystems Quant Studio 3. Two technical replicates were performed for each reaction and sample, and amplification of the correct products was verified by melt curve analysis. Relative gene expression was calculated with the following formula: Δ∁𝒕𝒕 = 𝟐𝟐^∁𝒕^ ^𝑹*p*132^/𝟐𝟐^∁𝒕^ ^16*s*^, ∁𝑡 are the mean ∁𝑡 value of the technical duplicates. Primer info and RT-qPCR conditions were listed in the supplementary Table S2 and Table S3.

### Survival assay

Flies were inoculated under CO_2_ anaesthesia via septic pin prick with 10 μm diameter stainless steel needles (Fine Science Tools, CA, USA). These needles were bent approximately 250 μm from the end to provide a depth stop and dipped in the inoculum before being pricked into the anepisternal cleft in the thorax of anaesthetised flies. This method has been demonstrated to be highly repeatable and gives a consistent initial dose (our estimate of initial dose (CFUs) introduced into individual flies for each pathogen: OD = 1 *P. rettgeri*: 10^(1.94 ± 0.08); OD = 1 *S. aureus*: 10^(2.75 ± 0.12)). All inoculations were done by a single experimenter (HS). Following pricking, the number of alive flies in each vial was recorded daily for 14 days. Systemic infection via wounding was used rather than oral infection to allow us to control pathogen dose. Flies were flipped into new food tubes (germ-reduced flies to sterile food under the flow hood, normal flies to normal food) every three days until the end of the experiments.

### Statistical analysis

Survival results were analysed using R v 4.3.1 (R core team, 2023) using a mixed effects Cox proportional hazards model [92]. A critical assumption of the Cox proportional hazards model is that when the predictor variables do not vary over time, the hazard ratio comparing any two observations is constant with respect to time [93]. The “cox.zph” function in the “survival” package [94] was used to diagnose whether the models fit the proportional hazards assumption of a Cox regression. The “strata” function was used to correct for the assumption of a Cox regression model. In this case, it is assumed that individuals in different strata have different baseline hazard functions, but all other predictor variables satisfy the proportional hazards assumption within each stratum. There were no estimations of the stratified variables [93, 94]. Interaction effects were tested using the anova function, model comparison and model simplification from the full model. Models included random effects of vial and experimental blocks to account for non-independence [95]. We observed that the hazard ratio estimates for microbiome effects remained stable both before and after correcting for the Cox regression model assumption using the strata function. The results presented are based on the model described below:

1. the full model with pathogen, species and microbiome treatment effect:

*coxme (Surv(time, status) ∼ Species*Pathogen + Species*Microbiome+ Pathogen*Microbiome+ (1|replicate/vial)*

1. the model at the individual species and pathogen level when the models fit the proportional hazards assumption:

*coxme (Surv(time, status) ∼ Pathogen +Microbiome + (1|replicate/vial)*

and the model at the individual species and pathogen level when the models do not fit the proportional hazards assumption:

*coxme (Surv(time, status) ∼ strata(Pathogen) + Microbiome + (1|replicate/vial)*

In model 1) Three way interactions were removed to avoid introduction of complex interaction term. In both model 1) and model 2), the control treatment (Ringer pricked flies) and the normal flies without manipulation of the host microbiome were set as reference.

All data and R code for analysis and data visualisation has been uploaded to an online repository https://figshare.com/s/0f14bc4ffba8b632eeca.

Because of a lapse in American government funding for NCBI, the sequencing data we submitted via the NCBI website have not been processed for a long time (Bio Project ID PRJNA1345360). We have uploaded the raw Amplicon sequencing data to the online repository https://figshare.com/s/0f14bc4ffba8b632eeca.

## Results

We inoculated a total of 5691 flies across 64 treatments (structured as full factorial design: four species x four microbiome treatments x four pathogen treatments) in eight biological replicates. The number of replicate vials for each treatment varies due to the variability of newly emerged number of flies (mean = 7, range 2-8, for *D. putrida* recolonized *Acetobacter* flies, the Ringer treatment includes two replicates, while most of the other treatments have more than five replicates, see Table S4). Vials with fewer than 6 flies were excluded from analysis, so that a total of 5580 flies were finally accounted in the analysis. The pathogen treatments included a control Ringer’s solution [96] or one of the three pathogens listed above. Each biological replicate (vial) contained a mean of 13 flies (range 6-20).

### Drosophilidae 16s microbial composition vary between species and treatments

We assessed the composition and relative abundance of bacteria for each fly species for each microbiome treatment (Figure 2), and for individual samples (Figure S2). Across all fly species with normal intact microbiomes, the dominant bacterial phyla were Proteobacteria, followed by Firmicutes, and then a small proportion of Bacteroidota and Actinobacteriota. The most abundant bacterial families are Acetobacteraceae, followed by Lactobacillaceae, consistent with previous work [56, 57, 59, 62]. At the Genus level, in normal flies, *Acetobacter* was the most dominant genera across all species. Besides *Acetobacter*, *D. melanogaster* has a high level of *Fructilactobacillus*; *D. putrida* contains a high proportion of *Commensalibacter* and *Enterococcus*, *Providencia spp* made up 2.7 % of the community in normal flies of this species. In *S. pattersoni,* the relative abundances of *Levilactobacillus and Morganella* were higher than *Acetobacter* on average but note the pattern varies slightly between samples (Figure 2, Figure S2). *Z. davidi* has a relatively simple bacterial community, of low alpha diversity, dominated by *Acetobacter* and *Fructilactobacillus* (Figure 2, Figure S2, Figure S4).

**Figure 2.**
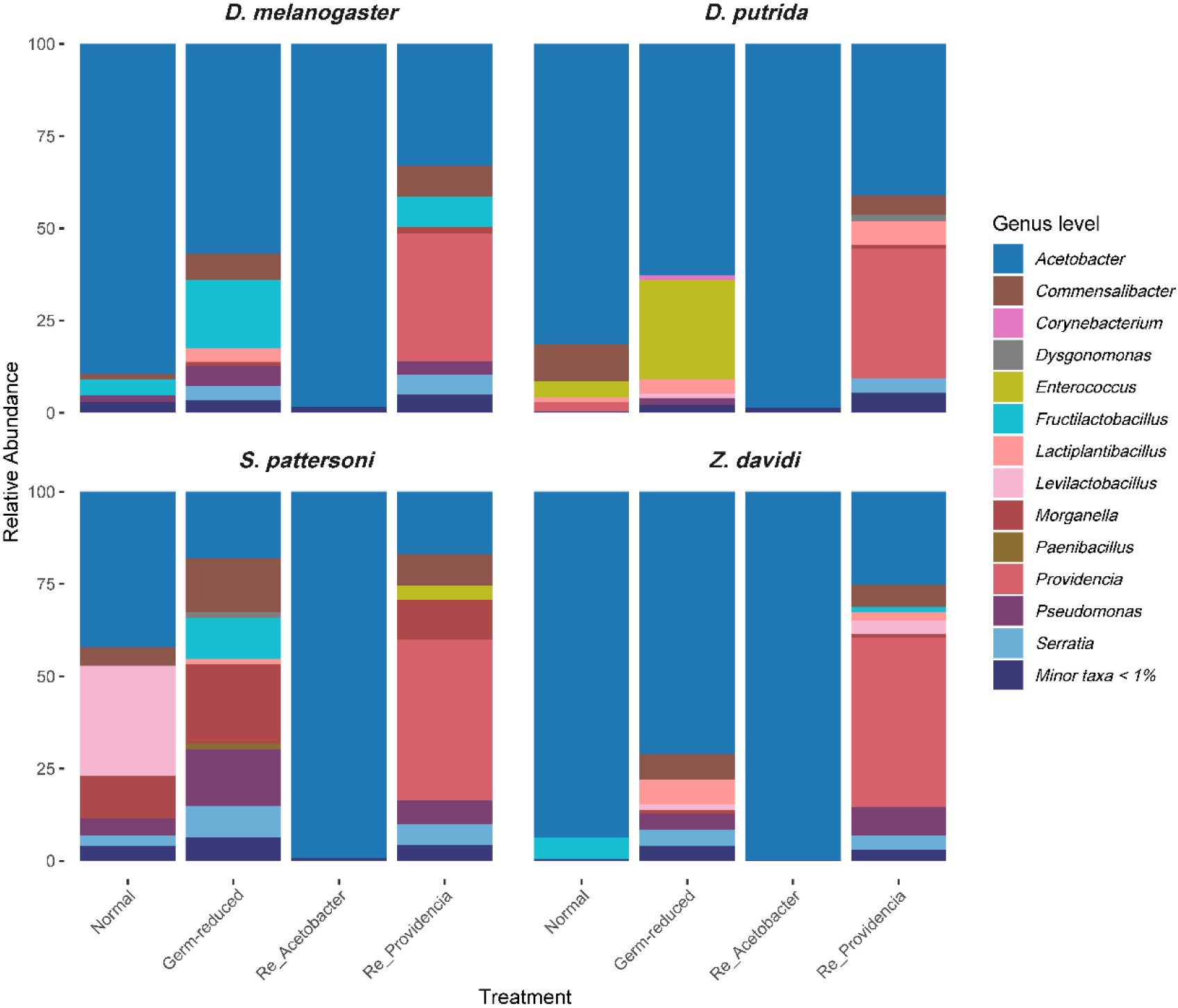
**Bacterial composition at the genus level, based on16S amplicons in each host species and microbiome treatment**. Plots are means across samples. Variation between individual samples were shown in Figure S2.

The microbiome reconstitution treatments had qualitatively different consequences: recolonisation rate with *Acetobacter* produced communities almost entirely dominated by this species, while recolonisation with *Providencia* produced more diverse communities where this genus constituted 35% - 46% of the community and relatively low microbiome load (Figure 2, Figure S2, Figure S3).

In germ-reduced flies the abundance of culturable microbiota were estimated at 10^4.04(±0.12)^ CFUs per fly lower than the normal flies (Figure S1). The relative 16s gene expression also indicated much lower abundance in this treatment (∼200 fold fewer) relative to normal flies (Figure S3). In germ- reduced flies the microbial composition was much more complex than the normal flies, indicating the disruption of microbiota by antibiotics dramatically inhibited the growth of dominant genera but did not fully clear the host microbiota (Figure 2, Figure S2). Germ-reduced flies exhibited a higher alpha diversity in bacterial reads compared to normal flies (Figure S4). This is likely because the reduction in bacterial numbers increased stochasticity and heterogeneity.

The DNA community standard control matched the quantitative ratio that provided by supplier (Figure S5), indicating no inherent biases in our sequencing and analytical approach. The microbial community standard (which underwent RNA extraction and cDNA synthesis) control showed variation in relative abundance of each genera, which likely reflects the actual active bacteria proportion in the sample (Figure S5).

### Susceptibility is shaped by host species, pathogen and host microbiome

Survival to different pathogens varied greatly across four tested species (Figure 3, host-by-pathogen interaction Chisq = 322.91, df = 9, p < 0.0001). For instance, DCV infections led to rapid death in two out of four cases killing ∼100% of *D. melanogaster* and 86% of *D. putrid*a flies within 14 days, but were less virulent to *S. patterson*i and *Z. davidi* (29% and 12% dead by day 14 post-infection), consistent with previous results [97]. The bacterium *P. rettgeri* killed all *Z. davidi* in three days post-infection but led to only 21% dead in 14 days in *D. putrida* (Figure 3). This pattern is consistent with previous results and confirmed that host species vary in their mortality in a pathogen-specific manner [73].

**Figure 3.**
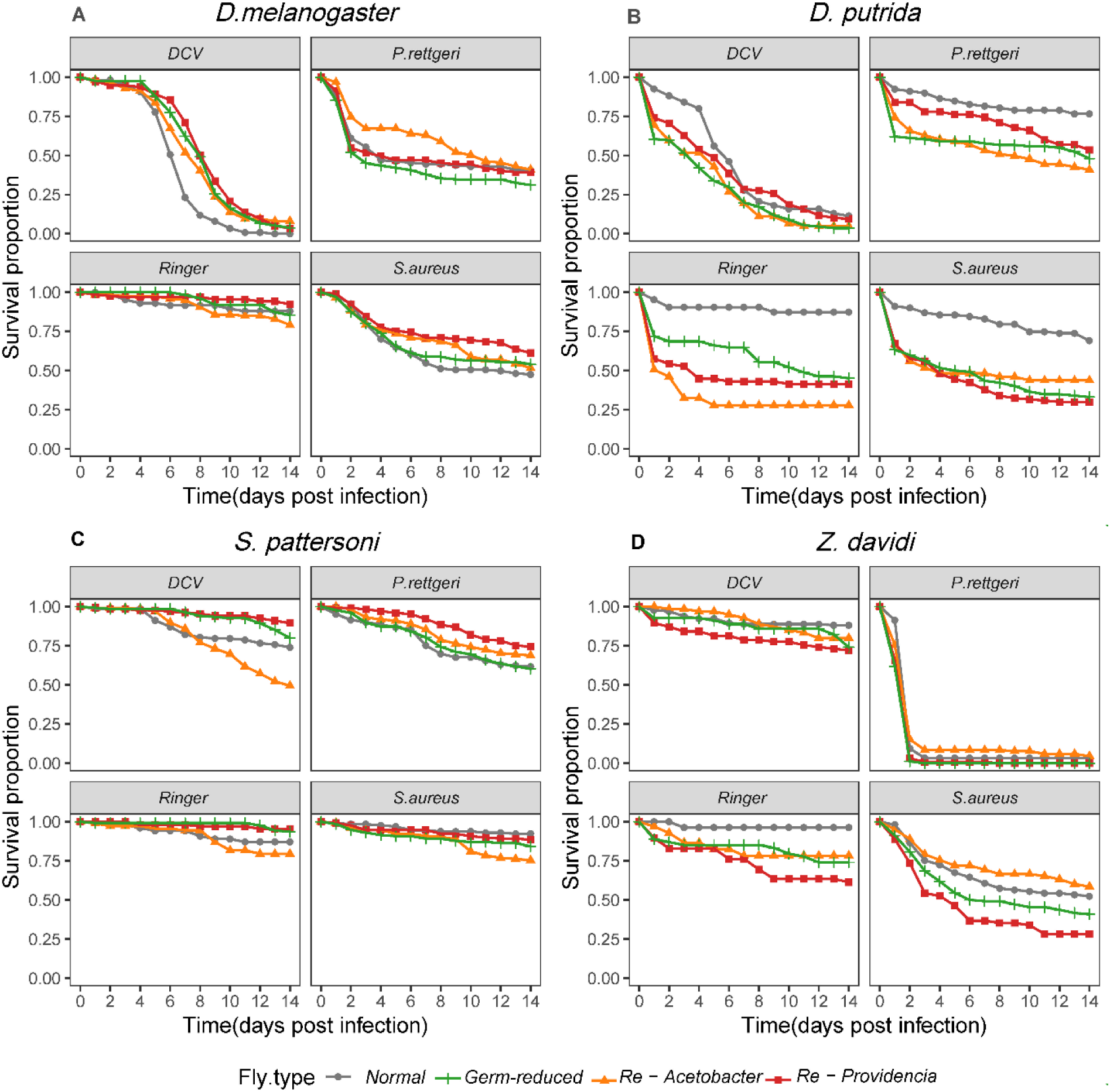
Survival to different pathogens across four Drosophilidae species. Survival curves show the survival proportions of four fly species inoculated with either: DCV, *P. rettgeri*, Ringer control *or S. aureus*. The X axis represents the time post-infection, and the Y axis represents the mean proportion of flies alive. Different colours represent either conventionally reared normal flies with intact microbiomes or germ-reduced flies or flies reassociated with *Acetobacter* (*Re- Acetobacter*) or *Providencia* bacteria (*Re – Providencia*).

Susceptibility was also driven by interactions between host species and microbiome treatments (Chisq = 75.51, df = 9, p = 0.0001), suggesting the microbiome effects vary greatly between different species. However, host susceptibility was only weakly affected by interactions between pathogen and microbiome treatments (Chisq = 19.86, df = 9, p = 0.02). The microbiome treatment had a strong interaction with host species, but not with pathogen type, suggesting that overall resilience to wounding could depend on the microbiome. This is evident for *D. putrida* in which survival rate following pinprick with Ringer’s solution varied strongly with microbiome treatment.

### Differing impacts of microbiome manipulation across host species

We then examined how normal flies, germ-reduced flies and flies with a reconstituted microbiome varied in susceptibility to different pathogen infection in each species. In *D. melanogaster*, we saw an improvement in survival following DCV infection in germ-reduced flies relative to normal flies (HR ± SE: 0.42 ± 0.27, p = 0.001), but no such effects were observed for bacterial infection (Figure 3A, Figure 4, Table S4). In DCV infection, treatments that manipulate the microbiome improved host survival in terms of the timing of mortality: normal flies with intact microbiomes died more rapidly (Figure 3A, Figure 4), but had similar final levels of total mortality.

**Figure 4.**
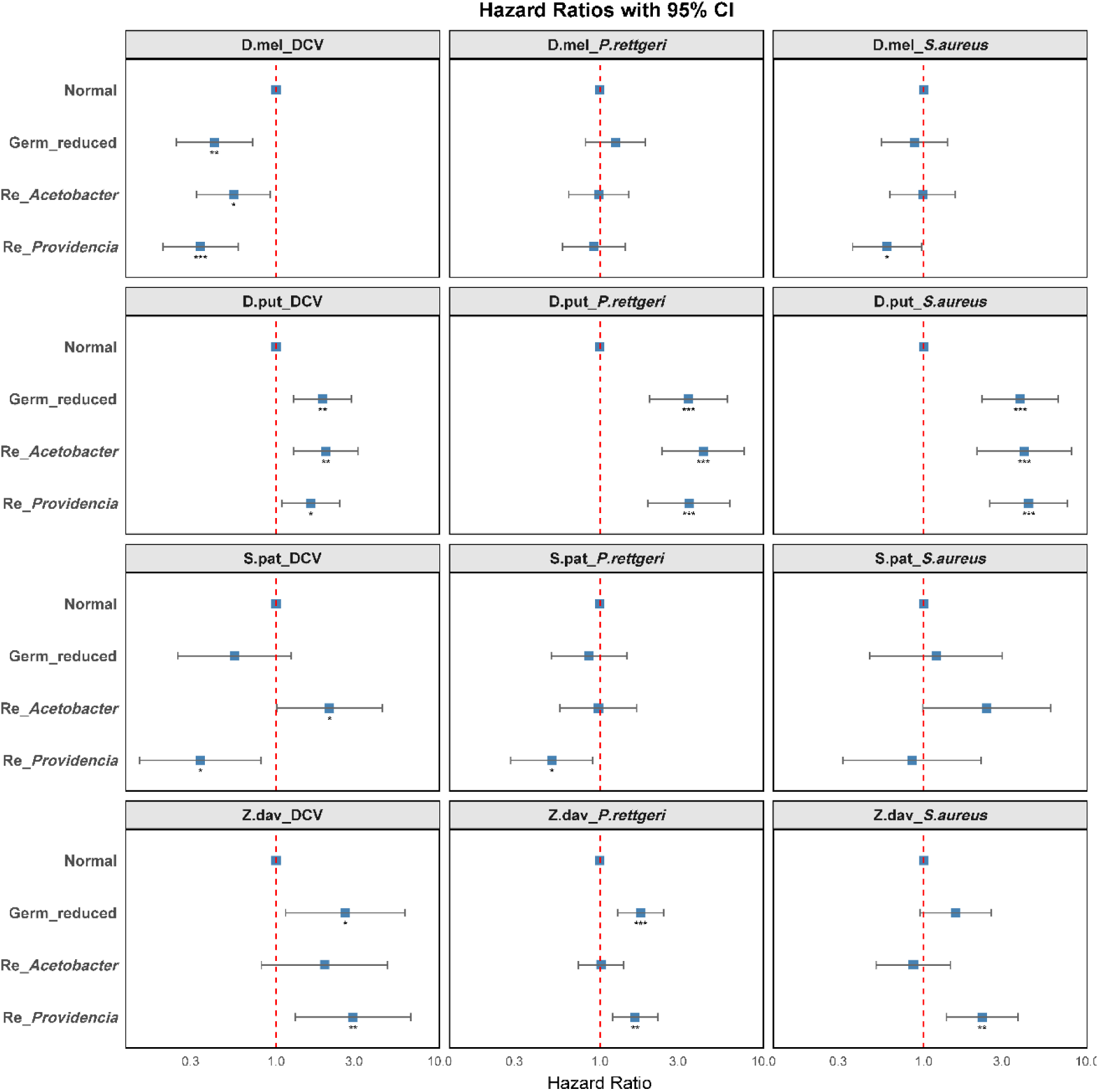
Hazard ratios across treatments. Plots show estimates of the hazard ratio for fixed effect terms of microbiome treatment in model 2 (pathogen effects excluded), which has accounted for the independence of random effects. The X axis represents the hazard ratio and 95% confidence interval in microbiome-manipulated flies relative to normal flies; a hazard ratio of <1 indicates increased survival relative to flies with normal microbiomes, a hazard ratio >1 indicates reduced survival relative to normal flies. Significance levels are highlighted as asterisks: * p < 0.05, ** p < 0.01, *** p < 0.001.

We observed strong effects of microbiome manipulation in *D. putrida* (Figure 3B); in both bacterial and DCV infection, germ-reduced *D*. *putrida* had reduced survival compared to flies with intact microbiomes (Figure 3B, Figure 4, Table S2). Flies recolonised with either *Acetobacter* or *Providencia* had reduced survival compared to normal flies (HR ± SE: *Re-Acetobacter*: 2.59 ± 0.17, p<0.001, *Re-Providencia*: 2.33 ± 0.16, p < 0.001). However, the hazard ratio estimate was close to 1 when using the germ-reduced flies as reference (HR ± SE: *Re-Acetobacter*: 1.02 ± 0.16, p = 0.88, *Re- Providencia*: 0.91 ± 0.14, p = 0.55), indicating that there were no statistically significant differences between germ-reduced flies and those with reconstituted microbiomes.

The survival of *S. pattersoni* was generally very high (Figure 3C). Disruption of the microbiome had no detectable impacts on the outcome of bacterial infections (Figure 4, Table S4). There were detectable microbiome effects on DCV infection in terms of the survival between normal flies and microbiota-reconstituted flies. Flies recolonised with *Acetobacter* had poorer survival than normal flies (HR ± SE: 2.12 ± 0.37, p = 0.04), but flies re-colonised with *Providencia* improved their survival compared with normal flies (HR ± SE: 0.34 ± 0.43, p = 0.01) (Figure 3C, Figure 4).

For *Z. davidi*, following DCV infection, normal flies survived better than germ-reduced flies (Figure 3D, Figure 4). Flies recolonised with *Acetobacter* had relatively higher survival than germ-reduced flies in both *P. rettgeri* (HR ± SE: 0.57± 0.16, p = 0.0007) and *S. aureus* (HR ± SE: 0.54 ± 0.23, p = 0.01) infection although we note that *P. rettgeri* infection caused extremely low survival. Flies recolonized with *Providencia* had reduced survival in most cases even in control treatments (Table S4).

## Discussion

Our study demonstrates that it is oversimplistic to assume that microbial symbionts have a protective role in terms of defending hosts from systemic infection. Microbiome components can increase, decrease or have no effect on host susceptibility in a way that depends on host species and the pathogen in question. Even within well-studied viral infections, we found that manipulation of the microbiome could either increase or decrease host mortality following pathogen challenge. Patterns were also not consistent across bacterial pathogens. It is important to point that this study did not randomly select host species: we deliberately included species (*D. putrida, S. pattersoni*) that we expected to show protective effects from the microbiome in order to explore the diversity of host-microbiome interactions. The strong effect of microbiome manipulation in *D. putrida* is therefore unlikely to be representative of the group. These results suggest that protection from systemic infection is not the prime selective force shaping microbiome composition in Drosophilidae. Other considerations such as lack of pathogenicity [37], nutrient acquisition [26, 58, 65] or trade-offs between control and costs [98] are potentially more important. While this study did specifically not consider interactions between the gut microbiome and intestinal pathogen, any beneficial effects of microbiota via immune priming may therefore be largely down to chance [99, 100].

### Biological underpinning of microbiome effects across tested species

In *D. putrida* we successfully reconstituted an *Acetobacter* dominated community (using an isolate from this species, see Figure 2). It appears that it is likely to be original disruption of the microbial community with antibiotics that is driving this change in survival rather than the single strain of microbiome manipulation. We suscept this for two reasons: i) we noticed there is large proportion of flies dead in Ringer infected control in both germ-reduced flies and microbiome manipulated flies specifically in this species (Figure 3B, day 14 Survival for Ringer inoculated normal flies: 86%, for Ringer inoculated germ-reduced flies: 49%), indicating the overall resilience to pinprick infection in *D. putrida* are generally low. ii) There were no survival differences between *recolonized Acetobacter* flies, *recolonized Providencia* flies compared to the germ-reduced flies, indicating the susceptibility change were caused by overall disruption of the microbial community rather than single microbiome strain. In addition, we also observed some mortality during the 3-day microbe recolonization period in *D. putrida* flies, suggesting that individual microbiome strains, when present in higher load, exhibit pathogenic effects in the host.

A straightforward immune priming interpretation of these systemic infection data is open to question. First, manipulating the microbiome had consequences for susceptibility to mock infection in *D. putrida* in particular. While mock infection might introduce a tiny dose of microbes into the haemolymph via the cuticle, an alternative explanation, such as interspecies variation in wound recovery, could be more relevant. Second, we successfully reconstituted two microbiome treatments using bacteria sourced from *D. putrida*: one dominated by *Acetobacter*, and another more diverse community but low abundance containing *Providencia*. This prompts the question as to why these communities did not offer improved survival similar/relative to the normal flies. Potentially, the less-abundant members of the *D. putrida* community are providing key functions here. More likely, the microbiome structure as a whole may be less important in systemic infection, the microbiome effects in certain species may indicate either species specificity or the just the side-effects after using antibiotics in certain species.

In *D. melanogaster*, we observed evidence that the native microbiota may facilitate DCV infection, as germ-reduced flies improved host survival. While microbiome manipulation delayed mortality, it did not change the final outcome. The inconsistent survival pattern of DCV infection between *D. melanogaster* and the other species may suggest that microbiome effects during systemic infection are limited and highly context-dependent. These effects appear insufficient to significantly alter infection outcomes, which may be primarily determined by pathogen virulence and host immune responses [101, 102].

In *S. pattersoni*, overall survival was high, which may have masked any microbiome effects during bacterial infection. However, recolonised *Acetobacter* flies of this species showed reduced survival, particularly during DCV infection. Since DCV can cause intestinal obstruction [103] and *Acetobacter* has been linked to abdominal bloating [104], their co-infection may exacerbate disease, an effect also noted in this study. Further experiments are needed to explore this interaction.

In *Z. davidi*, both germ-reduced and recolonized flies showed decreased survival across infections, a pattern similar to those observed in *D. putrida*. The dominated microbiome, which is *Acetobacter* and *Providencia*, may have chances to provide genuine protective effects through immune priming [69]. When germ-reduced flies are recolonised with *Providencia* or *Acetobacter* and then exposed to a novel pathogen, co-infection may occur, eliminating any protective benefit. These could possibly explain the reduced survival observed in both *Z. davidi* and *D. putrida*. Nevertheless, it is important to note that the effects of microbiome disruption are not entirely consistent between these two species.

### Effects of antibiotic treatment

Our results indicated a strong effect of microbiome manipulation in *D. putrida* for both mock infected and pathogen infected flies but only weak effects in other species. A parsimonious explanation is that disturbance of the microbiome with antibiotics early in adult life had a physiological impact on *D. putrida* that made it less robust. We chose to use RNA-based 16S amplicon sequencing, as RNA-derived community profiles are more likely to capture the active fraction of the microbiome, which may play a direct role in host metabolic and immune processes [83, 105]. We noticed the germ- reduced flies were not as “sterile” as we expected. This could reflect both technical limitations in sequencing axenic samples and the inherent difficulties of generating truly germ-free flies across species. Our antibiotic cocktail recipe was optimised by plating isolated microbiota to a range of antibiotic-containing media. While we sought to eliminate spurious effects of antibiotic carry-over at the point of infection, such as by transferring flies to sterile food for a few days before infection, we cannot entirely eliminate the physiological consequences of antibiotic dosing [106], which may have been considerable in *D. putrida.* We did not see any evidence of any consistent protective effect of antibiotic exposure, this would produce a pattern in which all germ-reduced flies would have increased survival on challenge with bacteria so we believe our protocols have eliminated any direct antimicrobial-microbiome interactions. However, we cannot exclude indirect consequences of antimicrobials on insect physiology. Work leading up to this project suggests that a shorter period between dosing of insects and pathogen challenge could have produced more controlled microbiomes, but with the trade-off of increased carryover of antimicrobials.

### Infection routes and ecological context

Here we infected our flies via wounding as this avoids variation in susceptibility driven by species-specific differences in feeding rate and inoculation dose. However, in doing so we bypassed gut immunity and a major point of contact between microbiota and pathogen. Many studies has shown the protective effects of gut microbiota either in defence of oral infections [48, 107] or improving the life span [43]. However, when examining the effects of removing/disrupting the gut microbiome and observing the general immune stimulation response in the host, it is always important to consider the ecological context in which these interactions occur. When microbiome and pathogen occupy the same compartment (e.g., the gut), the potential for direct and indirect interactions increases. For example, in *C. elegans*, *Lactobacillus. acidophilus* significantly decreased the burden of a subsequent *Enterococcus faecalis* infection in the intestine, and prolonged the survival of nematodes exposed to pathogenic strains of *E. faecalis* and *S. aureus* [51]. In lepidopteran insects such as *Galleria mellonella*, the *Enterococcus*-dominated gut microbiota in host protect against Gram-positive bacterial and fungal pathogens [40]. However, it is still unclear whether the gut-derived signals affect immunity elsewhere or if immune cells and metabolites can migrate to systemic sites, affecting the outcome of systemic infections.

### Host specificity and phylosymbiosis

Our earlier studies have indicated that both *P. rettgeri* and *S. aureus* infection across Drosophilidae species showed high phylogenetic signal in susceptibility. In *P. rettgeri* infections 94% of the variation in mortality can be explained by host phylogeny, and 64% for *S. aureus* [73]. These results suggest that some microbiome members that have co-evolved with their hosts, susceptibility to infection may be largely determined by host related factors. Millions of years of co-evolutionary divergence may have shaped infection outcomes across this family, leading to high specificity in certain lineages. For example, the survival of *S. pattersoni* was very high, suggesting broad and effective immunity in this species. *Z. davidi* was highly susceptible to bacterial infection especially following challenge with *P. rettgeri* but was tolerant of DCV infection [73]. This may indicate different immune pathway evolution in these species. The high phylogenetic signal could also be due to related species having more similar microbiomes, which could affect susceptibility positively or negatively [108]. However, co-diversification of the microbiome between related species does not mean co-evolution. Several species have shown the patterns of phylosymbiosis [5, 16–20, 109–113]. These does not necessarily imply the microbiome have coevolved with hosts, as the microbiome structure is dynamic; balanced and shaped by not only hosts but also diet and environmental factors [14, 114, 115]. Selection pressure on microbes to provide positive benefits to host may therefore be rare and/or intermittent [15].

It is important to recognise that many animal hosts tend to be relatively permissive in terms of microbial colonisation [116]. While simple environmental filtering mechanisms, for example, pH regulation, can help shape microbial communities at relatively low cost. Maintaining tight control over these communities through high expression of immune genes can be very costly. It will form selective pressure for the evolution of immune genes, driving the occurrence of polymorphisms of immune effectors to defend against specific bacterial infection [104]. Coevolution is more likely to emerge when microbes are transmitted vertically with high fidelity across generations [117]. Very high levels of community control usually require specialized anatomical features or ecological niche adaptions. These conditions are potentially not met for hosts that consume large numbers of microbes as part of their diet. In summary, caution is required when generalising about the potential beneficial impacts of microbiomes, particularly microbiome-mediated protective effects through mechanisms such as immune priming or host-microbiome co-evolution.

## Supplementary materials

**Figure S1** Bacterial load of normal flies and germ-reduced flies.

**Figure S2** 16S amplicon sequencing results showing bacterial composition in each host species and each type of flies at the genus level (individual sample).

**Figure S3** RT-qPCR results showing the fold change of 16s gene expression between different fly types.

**Figure S4** Alpha diversity between treatments.

**Figure S5** cDNA control showing consistent composition but inconsistent abundance compared with DNA control samples.

**Figure S6** Belta diversity between species and treatments.

**Table S1** Drosophilidae host species, and rearing diet information.

**Table S2** Primer information of RT-qPCR.

**Table S3** RT-qPCR cycle conditions.

**Table S4** Original survival data sheet.

## Acknowledgements

Thanks to Exeter sequencing centre for their help in 16s amplicon sequencing and the omics analysis. Thanks Dr Mark Hanson for his help in experimental design and the comments on an earlier draft of the manuscript.

## Open Access

**For the purpose of Open Access, the author has applied a CC BY public copyright licence to any Author Accepted Manuscript version arising from this submission.**

## Data and code availability

All data files and R scripts used in this study are available in an online repository: https://figshare.com/s/0f14bc4ffba8b632eeca

## Declaration of AI use

We have not used AI-assisted technologies in creating this article.

## Ethics

This work was assessed by the University of Exeter ethics committee (application no. 523751).

## Author Contributions

HS, BL, BR: Conceptualisation, Investigation, Methodology, project administration, Formal analysis, Writing- Manuscript draft, Writing- Reviewing and editing; HS: Formal Experiments, Data curation, Visualisation; BR, BL: Resources, Grant acquisition, Supervision.

