## Supplementary materials for "Effect of the microbiome on pathogen susceptibility across four Drosophilidae species"

**Supplementary material**

Hongbo Sun[1*], Ben Longdon[1+], Ben Raymond[1+]

1 Centre for Ecology & Conservation, Faculty of Environment, Science, and Economy, University of Exeter, Penryn Campus, Penryn, United Kingdom

* Corresponding author.

+ These authors contributed equally to this work.

**Author information:**

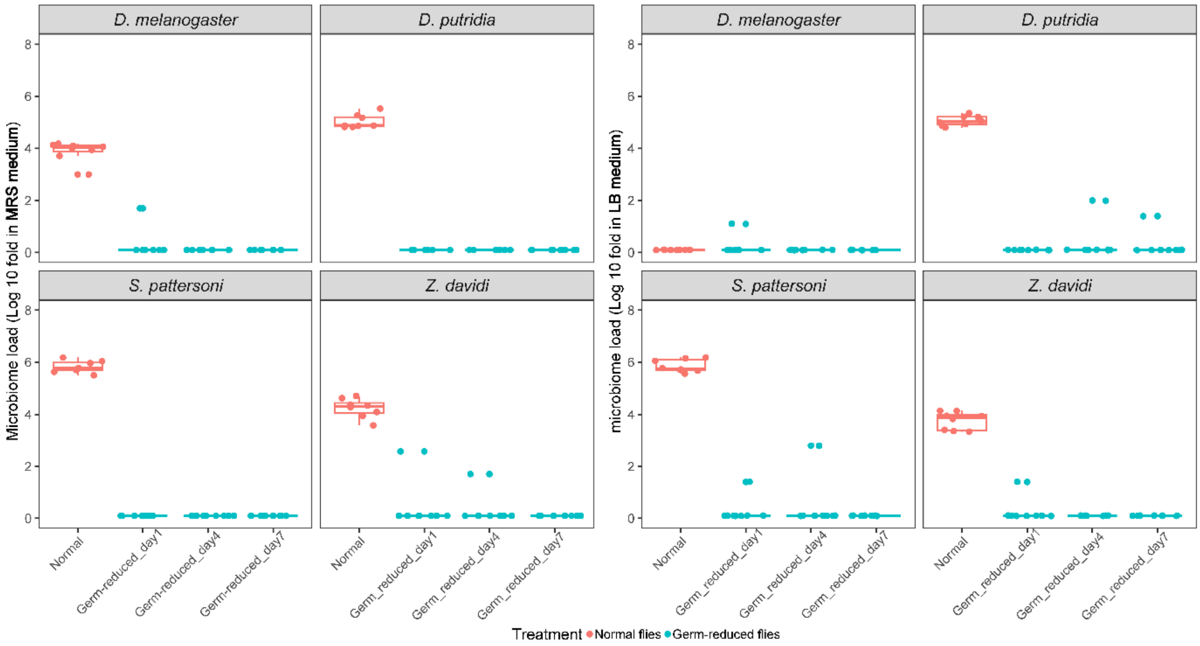


**Figure S1 Bacterial load of normal flies and germ-reduced flies.** Bacterial load (CFUs) of normal flies (red) and germ-reduced flies (blue) on selective media --- MRS (left) and LB (right) agar. Germ reduced flies were tested after one-week antibiotic food treatments, then CFUs were measured at three time points (day 1, day 4, and day 7) in the following week, each dot represents one individual fly. Each dot represents one individual fly collected from each treatment.

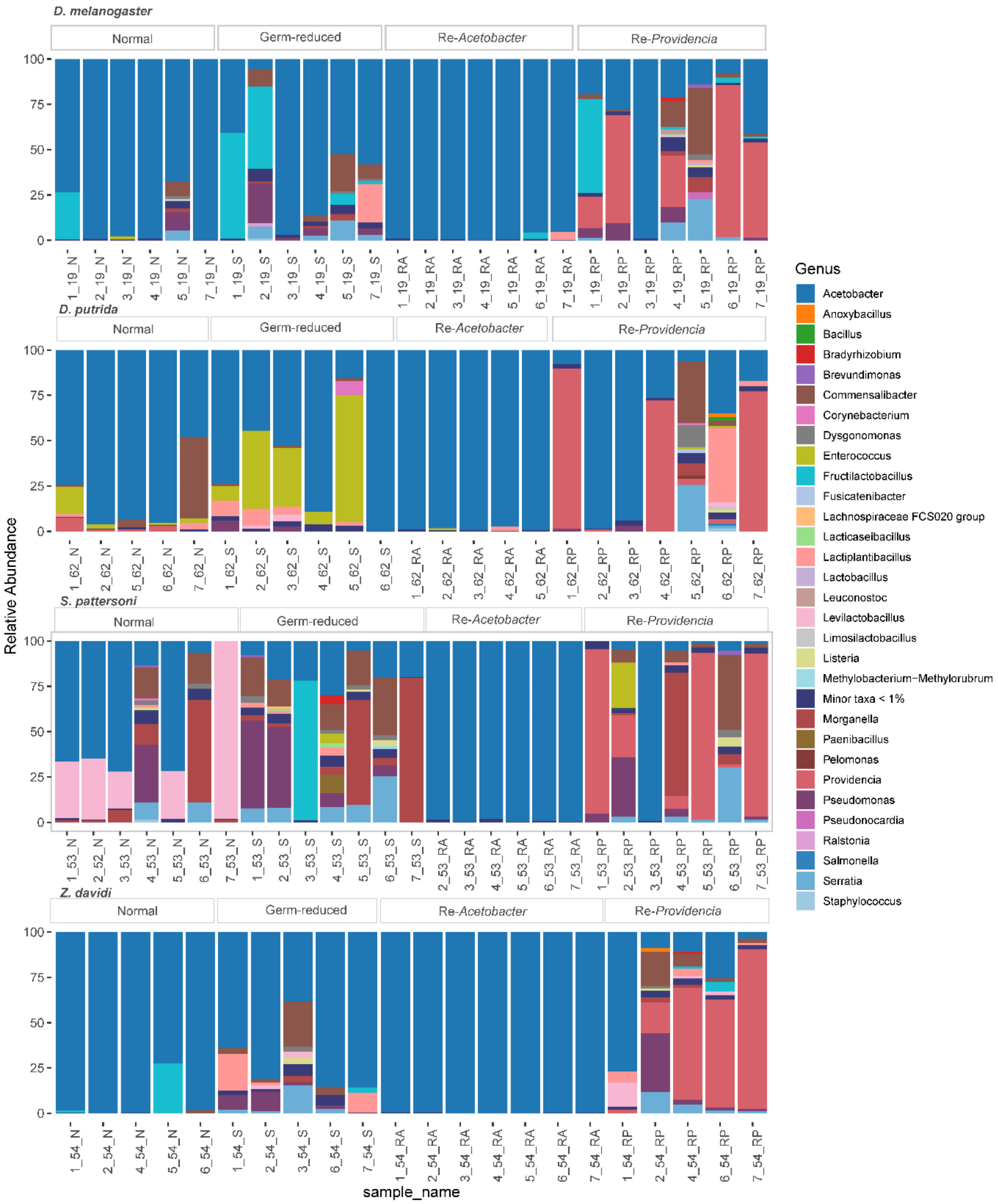


**Figure S2 16s amplicon sequencing results showing bacterial composition in each host species and each type of flies at genus level (individual sample).**

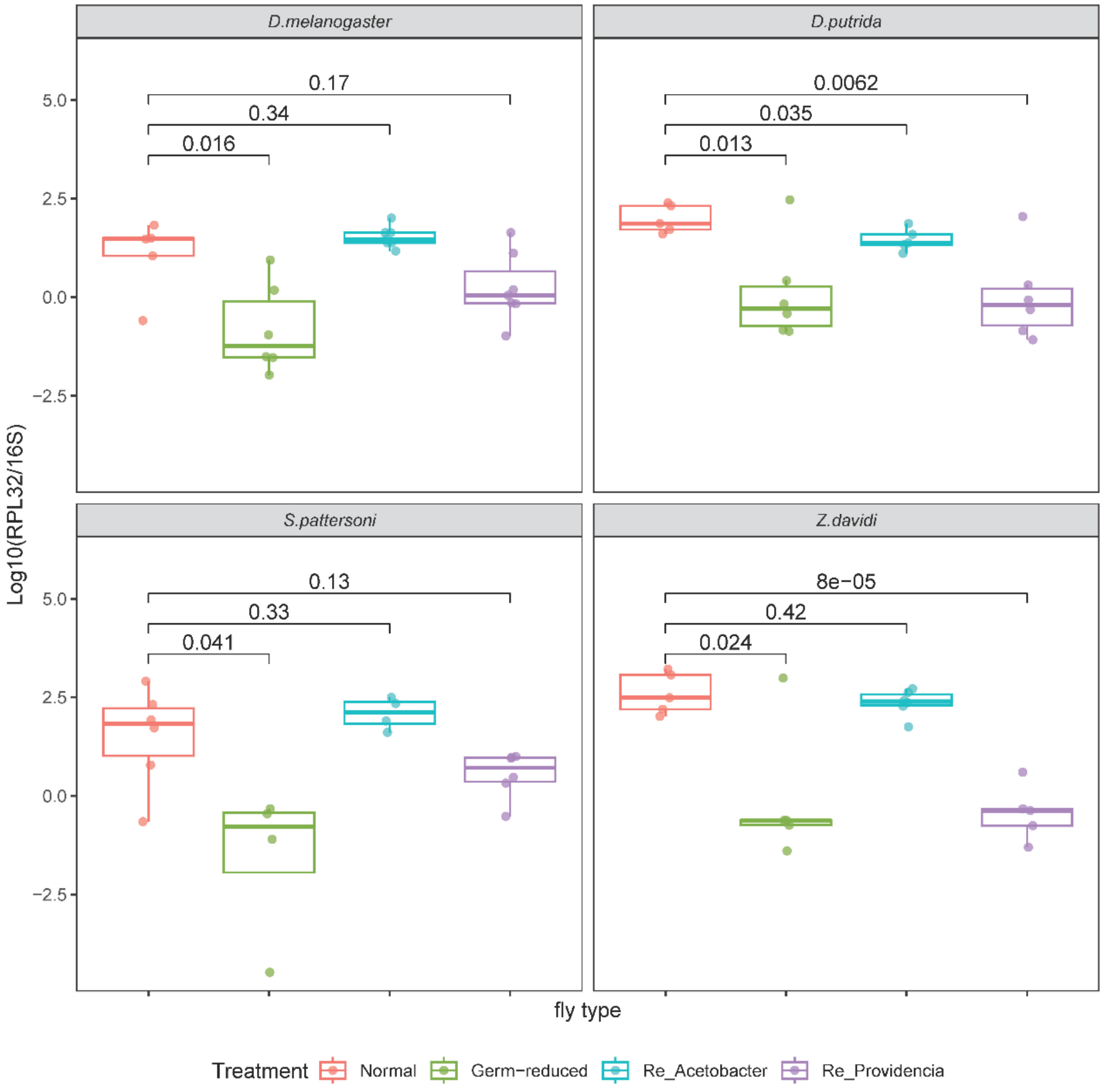


**Figure S3 RT-qPCR results showing the fold change of 16s gene expression between different fly types.** Each dot represents a pool of 8 flies from each block. p value were shown on top of the comparison bar.

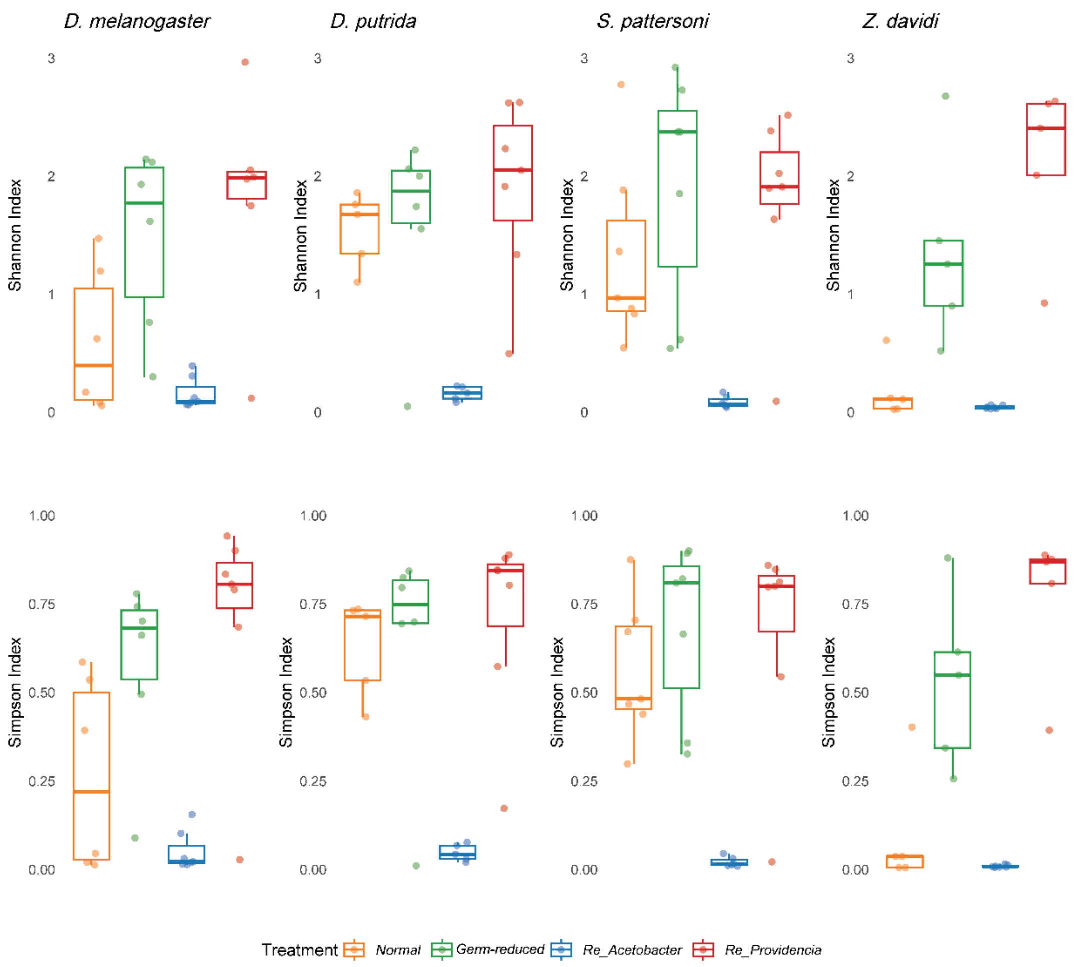


**Figure S4 Alpha diversity between treatments.** Alpha diversity between treatments showing Germ-reduced flies and Recolonized *Providencia* flies with relatively higher species diversity compared with those normal flies or flies recolonized with *Acetobacter*.

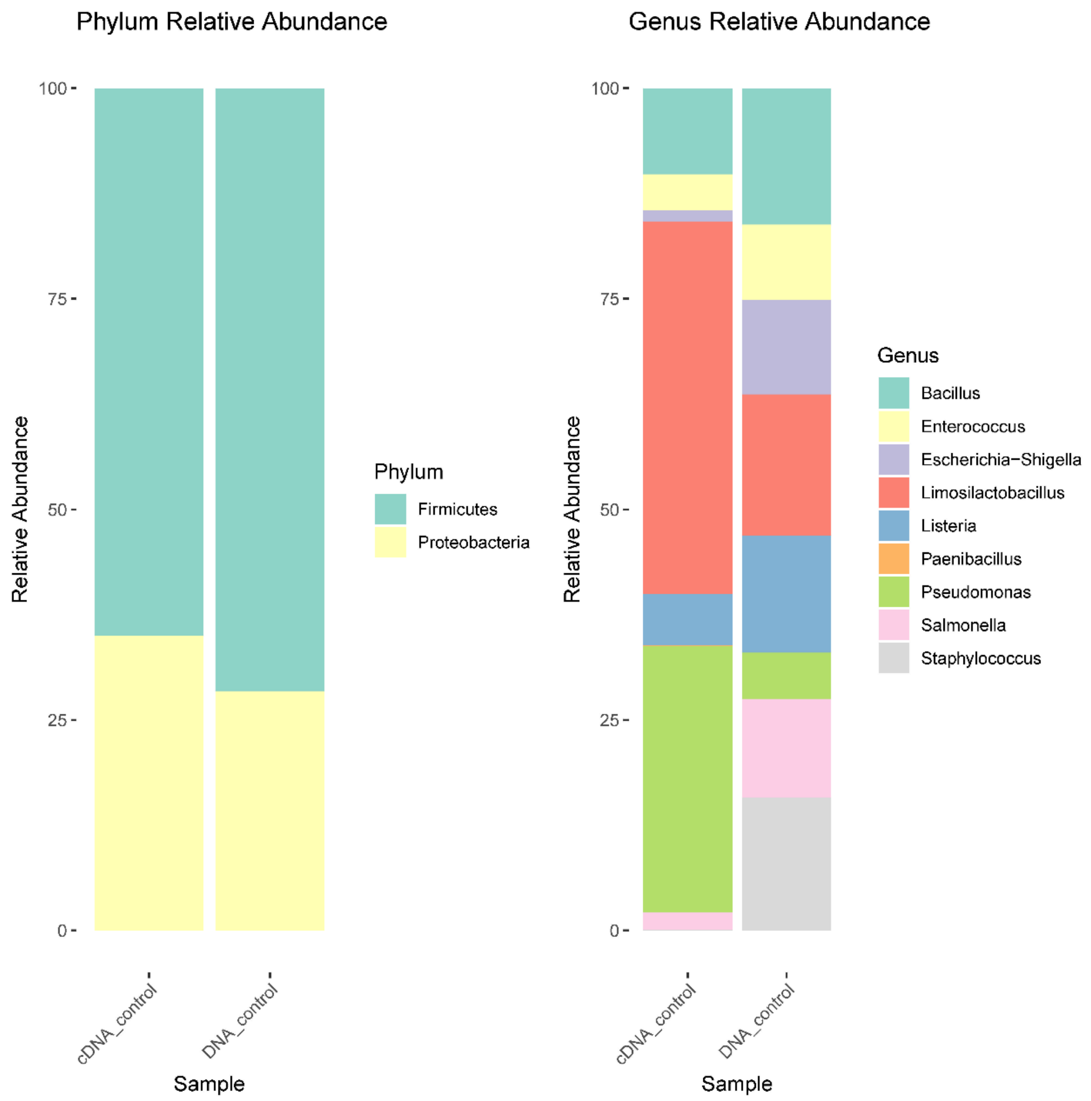


**Figure S5 cDNA control showing consistent composition but inconsistent relative abundance compared with DNA control samples.**

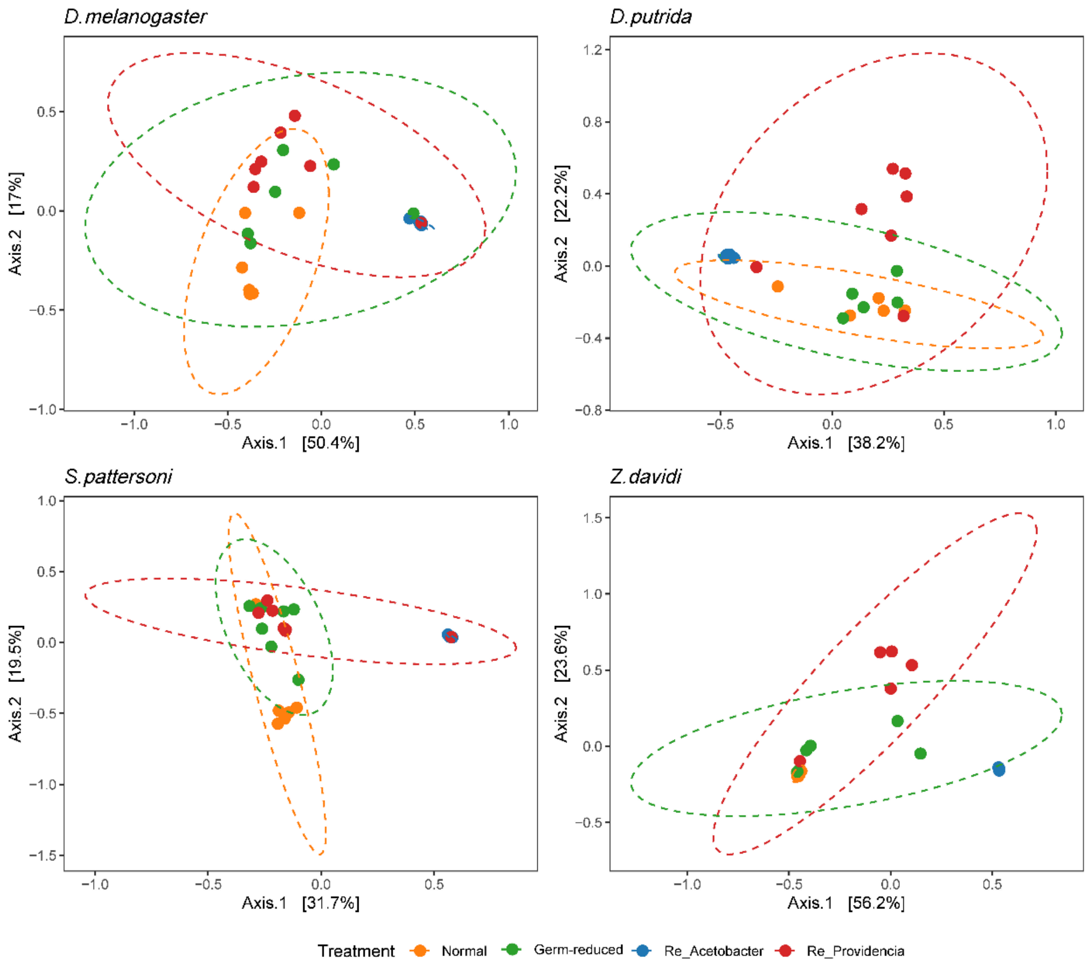


**Figure S6 Belta diversity between species and treatments.** Belta diversity between species and treatments showing all types of flies the species composition overlapped together.

| ID | Animal | Diet | Genus |
|---|---|---|---|
| **Before eclosion** | | | |
| 1 | *D. melanogaster* | cornmeal | Drosophila |
| 2 | *D. putrida* | propionic | Drosophila |
| 3 | *S. pattersoni* | banana | *Scaptodrosophila* |
| 4 | *Z. davidi* | banana | *Zaprionous* |
| **After eclosion and in formal experiments** | | | |
| 1 | *D. melanogaster* | malt | Drosophila |
| 2 | *D. putrida* | malt | Drosophila |
| 3 | *S. pattersoni* | malt | *Scaptodrosophila* |
| 4 | *Z. davidi* | malt | *Zaprionous* |

**Table S1: Drosophilidae host species, and rearing diet information.** Rearing food recipes can be found at https://doi.org/10.6084/m9.figshare.21590724.v1 Germ reduced flies, microbiome manipulated flies were reared on autoclaved malt food.

| Species | Primer | Target gene | Sequence |
|---|---|---|---|
| *D. melanogaster* | F-d | Rpl32 | TGCTAAGCTGTCGCACAAATGG |
| | R-h | Rpl32 | TGCGCTTGTTCGATCCGTAAC |
| *D. putrida* | F-d | Rpl32 | TGCTAAGCTGTCGCACAAATGG |
| | R-q | Rpl32 | TGAGCTTGTTTGATCCGTAAC |
| *S. pattersoni* | F-a | Rpl32 | TGCCAAGTTGTCGCACAAATGG |
| | R-m | Rpl32 | TACGCTTGTTGGAGCCGTAAC |
| *Z. davidi* | F-a | Rpl32 | TGCCAAGTTGTCGCACAAATGG |
| | R-c | Rpl32 | TGCGCTTGTTGGAACCATAAC |
| All species listed above | 16SUnivF | 16S | AGGATTAGATACCCTGGTAGTCC |
| | 16SUnivR | 16S | YCGTACTCCCCAGGCGG |

**Table S2:** Primer information of RT-qPCR.

| Step | Temp | Time | Cycle |
|---|---|---|---|
| Initial denaturation | 95˚C | 2 min | 1 |
| Denaturation | 95˚C | 5 sec | 40 |
| Annealing/Extension | 60˚C | 15 sec | |
| Melt curve | 95˚C-60˚C | 0.1˚C/Sec | 1 |

**Table S3:** RT-qPCR cycle conditions.
